# Functional Implications of Disordered Terminal Regions of *Macrotyloma uniflorum* Bowman-Birk Inhibitors: A Molecular Dynamics Study

**DOI:** 10.1101/125807

**Authors:** Abhishek Acharya

## Abstract

Bowman-Birk Inhibitors (BBI) – a class of serine protease inhibitors is of considerable interest due to their anti-inflammatory and anti-carcinogenic properties. Recent efforts have focused on understanding the structure and dynamics of these inhibitors, and the molecular mechanism behind its bioactive properties. BBI derived from Horsegram seeds is an interesting member of the class that exists as a number of isoforms that differ in length at the C- and N-terminal disordered regions. Interestingly, the length (or conversely, truncation) of the terminal regions affect whether the protein exists as a dimer or monomer. Here, we have investigated the mechanism of dimerization in Horsegram BBI. A recent study has proposed that the dimerization occur via a C-terminal hook that forms a salt bridge with the opposite monomer and is pivotal to the dimerization process. We have employed long computational simulation methods to predict the stability of the proposed C-terminal hook; we show that the terminal regions are highly disordered and the salt bridges are significantly solvent exposed. Further, using Hamiltonian replica exchange method, we have sought to obtain the conformational ensemble of the disordered terminal regions and have identified a conformational state that provides an interaction hot-spot that aids in the dimerization of HGI. Our analysis predicts an alternate model of dimerization that largely agrees with previous experimental studies and yet again, highlights the importance of intrinsically disordered region in tailoring the protein function.

## Introduction

Protease Inhibitors (PI) are a common occurrence in plants. Apart from occurring in storage tissues such as seeds and tubers, these proteins may also be induced in aerial plant tissues as a result of injury. They play a prominent role of in plant’s natural defense mechanism against insects and microbial infestation.^1,2^ The ability to interfere with the peptide hydrolysis reaction catalyzed by proteases involved in digestion are responsible for their anti-nutritional activity. A number of plant PI’s have been identified that are classified into various families based on sequence and structural homology.^3–7^ One of the earliest discovered were the Bowman-Birk Inhibitors (BBI) in *Glycine max*^8,9^, which have thereafter been identified in other plants belonging to families such as *Fabaceae* and *Poaceae*.^10^ These inhibitors have received considerable attention for their anti-carcinogenic properties in past decade, spurring research to understand the structural features and their functional implications.^11,12^ More recently, BBI from soybean has been shown to inhibit HIV-replication in macrophages.^13^ It is expected that a detailed understanding of the structure-function relationship of these enigmatic molecules would be immensely valuable in designing new anti-cancer therapies.^14^

BBI are inhibitors of trypsin and chymotrypsin type serine proteases. They are small proteins of masses 6 to 9 kDa, characterized by two homologous domains that together form a structure stabilized by seven conserved disulfide bridges. Each domain (SCOP classification: all-β) is composed of three anti-parallel β-strands and a protease-binding site.^15^ The two distinct binding sites, Anti-Trypsin-Loop (ATL) and Anti-Chymotrypsin Loop (ACL), afford the protein the ability to independently as well simultaneously bind and inhibit two proteases in a 1:1:1 ternary complex, apart from forming 1:1 complexes with trypsin/chymotrypsin.^16^ These exposed sites act as substrates for the proteases that typically recognize Lysine (on ATL) or Phenylalanine (on ACL) for cleavage, but rather form a complex that renders the proteases unavailable for further peptide hydrolysis.^17^ In monocots however, BBI’s have only a single reactive site for protease inhibition. Additionally, a larger group of BBI’s of mass 16 kDa exists, suggested to have formed as a result of gene duplication after the divergence of monocots and dicots.^18,19^ In many dicots, the 8kDa double-headed BBI form highly stable dimers which appear as 16 kDa species in gel filtration and SDS-PAGE.^20–22^ Particularly, dry Horse Gram (*Macrotyloma uniflorum/Dolichos biflorus*) seeds have four characterized isoforms of BBI (HGI-I to HGI-IV). With almost identical sequences differing only at the N-terminal region, all of them exist as dimers in physiological conditions (Table 1).^22^ Interestingly, in germinated Horse Gram seeds, these inhibitors exist as monomeric isoforms which is attributed to the proteolytic truncation of the N and C-terminal ends of HGI.^23^ Three different Horse Gram Germinated Inhibitors (HGGI-I, II and III) have been identified and characterized (Table 1). The differences in native state (monomer/dimer) and trypsin /chymotrypsin inhibitory constants among the isoforms necessitated an inquiry into the role of the terminal regions and the involved mechanisms.

**Table 1.** Horse gram BBI forms. # Data reproduced from Sreerama et.al. (1997).^22^ *Data reproduced from Kumar et.al.(2002)^23^.

| Horse Gram BBI Isoforms | N-terminal sequence | C-terminal sequence | Quaternary State | Trypsin Inhibitory Constant (K<sub>i</sub>s) | Chymotrypsin Inhibitory Constant (K<sub>i</sub>s) | pI Value |
| --- | --- | --- | --- | --- | --- | --- |
| #HGI-I | STDEPSESSK | PCKSSHDD | Dimer | $1.38 \times 10^{-7}$ M | $3.10 \times 10^{-7}$ M | 4.60 |
| #HGI-II | HHESTDEPSESSK | PCKSSHDD | Dimer | $1.00 \times 10^{-7}$ M | $0.15 \times 10^{-7}$ M | 4.80 |
| #HGI-III | DHHQSTDEPSESSK | PCKSSHDD | Dimer | $0.87 \times 10^{-7}$ M | $3.90 \times 10^{-7}$ M | 5.10 |
| #HGI-IV | DHHESTDEPSESSK | PCKSSHDD | Dimer | $1.20 \times 10^{-7}$ M | $4.60 \times 10^{-7}$ M | 5.60 |
| *HGGI-I | DEPSESSK | PCKS | Monomer | $1.98 \times 10^{-7}$ M | $4.77 \times 10^{-7}$ M | 4.99 |
| *HGGI-II | EPSESSK | PCKS | Monomer | $1.89 \times 10^{-7}$ M | $3.64 \times 10^{-7}$ M | 5.09 |
| *HGGI-III | SK | PCKS | Monomer | $0.98 \times 10^{-7}$ M | $7.73 \times 10^{-7}$ M | 5.28 |

Recent efforts to elucidate the mechanism of stable dimerization of HGI-III have revealed the role of the C-terminal Asp76^a^ residues and Lys24^b^ in stabilization of the dimer, possibly via salt bridge interactions.^24^ For clarity, residues coming from respective subunits will be indicated with superscript ^a^ or ^b^ wherever necessary. Kumar et. al. have recently proposed a model of dimerization wherein the C-terminal region forms a hook, stabilized by a salt bridge interaction between Asp76^a^-Lys71^a^ (Figure 1)^24^. This forces Asp75^a^ to adopt an orientation that facilitates association with Lys24^b^ to form an additional salt-bridge which holds the dimer. This structure is formed at both ends of the dimer in a symmetrical manner. Incidentally, Lys24 is also part of the anti-trypsin loop, necessitating dimer disruption for trypsin binding. The same was suggested by analysis of models, fluorescence quenching and size exclusion chromatography.^24^ Although the model of dimerization and experimental findings are significant, the mechanism of dimer to monomer transition is unknown. Also, the function of N-terminal region remains unexplained. For instance, dimerization mechanism of HGI involving C-terminal hook does not explain the role of the first six N-terminal residues and their simultaneous loss in HGGI forms along with the C-terminal region. We have sought to computationally probe HGI dimerization and possible mechanism of dimer to monomer transition. Previous computational studies have been limited to model analysis and short simulations (5-10 ns)^22,24^ which have provided limited insight into protein dynamics, especially that of the flexible terminal segments.

**Figure 1:**
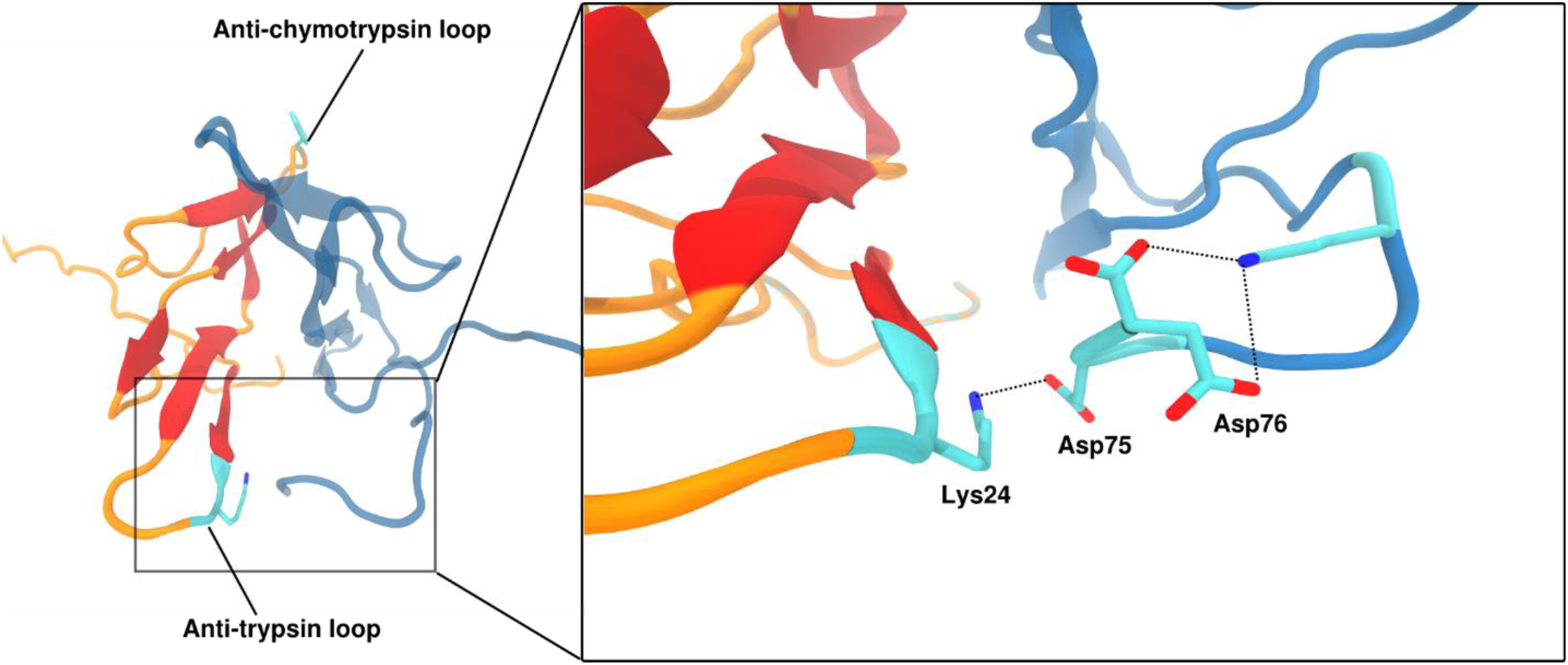
Model of the mechanism of HGI-III dimer formation involving a C-terminal hook. The dimer structure having two HGI-III monomers with the C-terminal in a hook like conformation. The Anti-Trypsin loop containing the Lys24 residue and the C-terminal residues Asp75, Asp76 and Lys71 are depicted in Cyan. The stabilization of C-terminal in a hook like conformation requires Asp76-Lys71 SB interaction, which enables stable SB between Lys24 and Asp75.

In the present work, we have performed computational studies on HGI-III dimer to identify for the first time a structural motif that may be critical for dimerization. We have undertaken long Molecular Dynamics calculations to analyze the general stability and dynamics of the inhibitor dimer and to identify prominent interactions that are involved in dimerization. Using REST2^25^ sampling method, we have attempted to identify most probable structures of the flexible N and C terminals and based on the analysis proposed a model to explain the existence of HGI-III as a dimer. Results obtained and conclusion drawn have been compared with previous experimental results wherever possible. The model supports the involvement of both N and C terminals in formation of a structural motif which provides an interaction “hot-spot” that may be crucial to stabilization of the dimer interface. Finally, we have suggested a possible mechanism of dimer to monomer transition which is essential for trypsin/chymotrypsin inhibition by HGI-III. Overall, the proposed model brings into perspective the structural significance of the terminal regions of HGI forms and its functional implications.

## Materials and Methods

### Sequence and Structural analysis

Sequences of BBI were retrieved from Uniprot Knowledgebase (http://www.uniprot.org/uniprot/). The redundant sequences were removed using CD-HIT clustering program using a sequence identity cutoff of 0.9. Multiple sequence alignment was performed using ClustalX program.^26,27^ The model of HGI-III used in simulations was obtained partly by homology modeling using Modeller9.14.^28^ As in previous modeling studies on HGI, ^21,24^ pea BBI crystal structure in dimer form (PDB: 1PBI) was used as a template with a sequence identity of 70.1%. The C-terminal region was modeled based on the recently proposed C-terminal hook model.^24^ Validation of the highest scoring model was done using PROCHECK.^29^ The truncated HGI models were prepared by manually editing the full length HGI model and energy-minimized. Surface Electrostatics were calculated by the Automated Poisson-Boltzmann Solver (APBS).^30^ Structures and MD trajectories were visually analyzed and rendered for figures using Chimera and VMD.^31–33^ Plots were prepared and rendered using Grace plotting tool (http://plasma-gate.weizmann.ac.il/Grace/).

## Computational Methods

For initial classical MD simulations of HGI-III and its truncated forms, the protein systems were prepared by placing the proteins in a triclinic box with 1.0 nm distance between box edges and protein in all directions. Proteins models were assigned parameters using the AMBER ff99SB-ILDN.^34^ The box was solvated using TIP3P^35^ explicit water model and system charges were neutralized by adding appropriate Na^+^ and Cl^-^ counterions. All systems were energy-minimized and equilibrated in NVT ensemble for 1 ns to gradually heat the system to 300 K and relax water molecules around the harmonically restrained protein. A restraining force of 1000 kJ mol^-1^ nm^-1^ was used for position restraints. The systems were next equilibrated for 1 ns with temperature and pressure coupling at 300 K and 1 bar. For initial temperature and pressure coupling, Berendsen weak coupling algorithm^36^ was used. Final production runs were conducted for 100 ns in the NPT ensemble. For the production runs, temperature and pressure was controlled using Nose-Hoover^37,38^ thermostat and Parrinello-Rahman barostat^39,40^ respectively. Non-bonded interactions were calculated using a cut-off of 1.2 nm for short-range interactions. PME (Particle Mesh Ewald) was used for treating long range interactions.^41^ All covalent bonds were constrained with P-LINCS algorithm.^42^ Water molecules were constrained to their reference geometry using SETTLE.^43^ The equations of motion were solved using the Leap-frog algorithm^44^ with a time-step of 2 fs. For each system, three simulations were conducted with randomly chosen seeds for generating atomic velocities which ensured independent runs. Gromacs 4.6.7^45,46^ was used for all MD simulations and analysis.

For obtaining canonical distributions of HGI-III and HGGI-I structures, REST2^25^ method was employed using Gromacs4.6.7 patched with modified Plumed 2.2.1. REST2 was implemented in Gromacs through this patch as described elsewhere.^47^ This implementation scheme allows individual replicas to evolve using different Hamiltonian. Specifically, in REST2 individual replicas are simulated at different effective temperatures using scaled potentials. The advantage of this method is that it allows the system to visit all relevant configurations at a much lesser computational cost compared to the conventional REMD method. The number of replicas required for REST2 scales as the square root of number of solute particles only as the water-water interaction energy terms do not have a bearing on the exchange acceptance rates.^25^ For the present work, we used 32 replicas with effective temperatures of the solute ranging from 300 to 600 K (λ from 1.0 to 0.5). We used AMBER ff99SB-ILDN^34^ for modeling proteins and TIP3P explicit water model. The simulations were run in NVT ensemble using Stochastic Velocity Rescaling^48^ for temperature control. Exchanges among replicas were attempted every 1000 steps with a time-step of 2 fs. Each of the 32 replicas for HGI-III was simulated up to 50 ns for a total of 1600 ns of simulations. For HGGI-I, a total of 800 ns (25 ns/replica) of simulations was performed. Each protein was energy-minimized and equilibrated for 1 ns in NVT ensemble before applying REST2 method. Cluster Analysis of the REST2 trajectories was performed using g_cluster utility from Gromacs 4.6.7 package. A clustering cut-off of 0.25 nm and 0.15 nm was employed for HGI-III and HGGI-I respectively. Clustering was performed using the gromos clustering method.

For the final 50 ns MD simulation of HGI configuration obtained from REST2 calculations, same method was employed as initial MD calculations.

## Results and Discussion

### Molecular Dynamics of HGI

In order to shed light on the dynamics of the N and C terminal regions and the stability of the salt bridges implicated in dimerization of HGI-III, simulations were conducted on HGI-III (HGI-1_76-FL) and its mutants (HGI_1-72, HGI_7-76 and HGI_7-72). The backbone RMSD plots (Figure 2, blue plots) and the average backbone RMSD values in Table 2 depict a general trend of large scale deviation from the initial configuration. This is due to the highly flexible 15 residues in the N – terminal and to a lesser extent the 6 C terminal residues that eventually converge to a more stable configuration (Figure 3). Specifically, the HGI variants with the full length N-terminal 15 amino acid peptide stretch (HGI_1-76-FL and HGI_1-72) show greater RMSD compared to proteins with a truncated N-terminal (HGI_7-76 and HGGI_7-72) (Table 2). In comparison, the backbone RMSD values of the core residues (16-70) have an average RMSD value of ∼ 2 Å (Figure 2, red plots). This indicates greater stability of the core 16-70 residues compared to the terminal regions.

**Fig 2.**
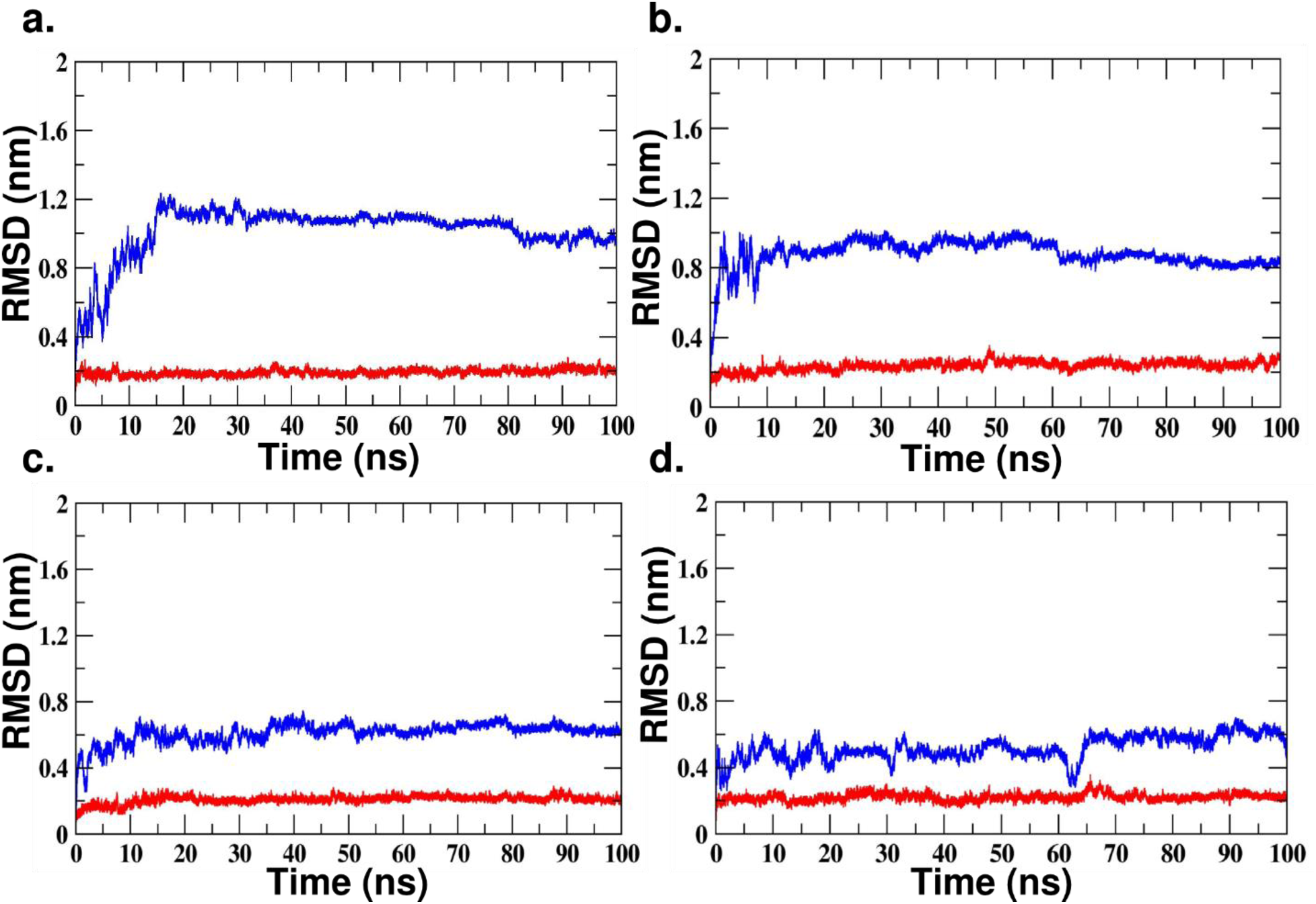
Backbone RMSD plots in case of a) HGI_1-76-FL b) HGI-1_72, c) HGI-7_76 and d) HGI-7_72. Blue plot depicts the backbone RMSD value of all residues in the simulated system. Red plots presents the RMSD calculations done only on the core residues from 16-70.

**Table 2.**
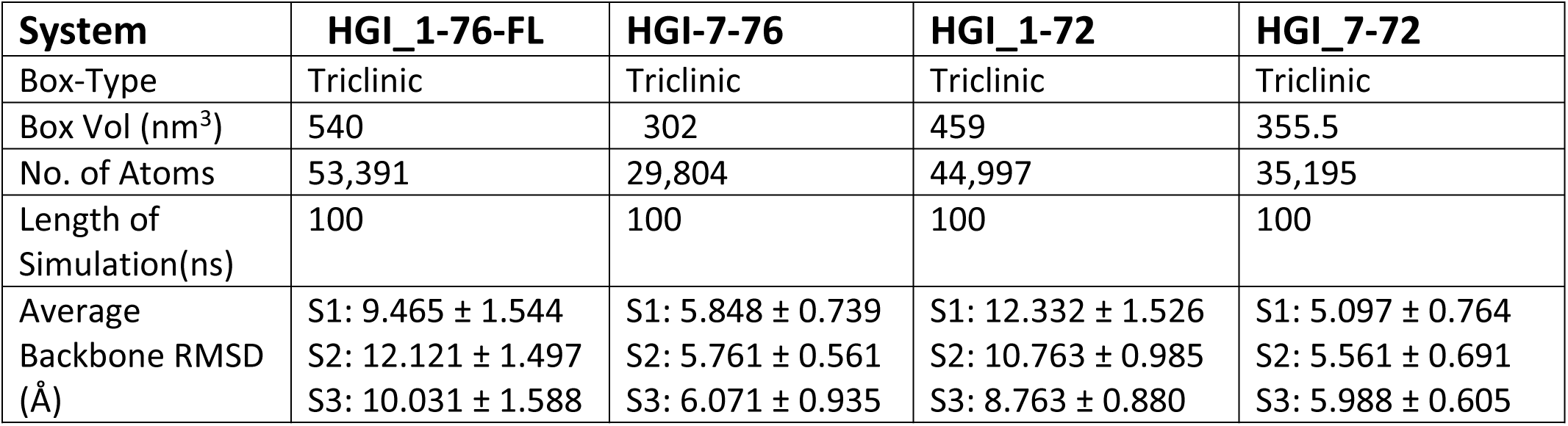
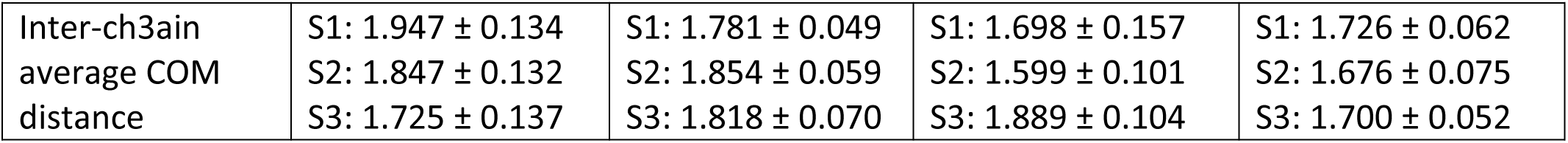
Classical MD simulation of HGI-III and its deletion mutants.

| System | HGI_1-76-FL | HGI-7-76 | HGI_1-72 | HGI_7-72 |
| --- | --- | --- | --- | --- |
| Box-Type | Triclinic | Triclinic | Triclinic | Triclinic |
| Box Vol (nm<sup>3</sup>) | 540 | 302 | 459 | 355.5 |
| No. of Atoms | 53,391 | 29,804 | 44,997 | 35,195 |
| Length of Simulation(ns) | 100 | 100 | 100 | 100 |
| Average Backbone RMSD (Å) | S1: 9.465 ± 1.544<br>S2: 12.121 ± 1.497<br>S3: 10.031 ± 1.588 | S1: 5.848 ± 0.739<br>S2: 5.761 ± 0.561<br>S3: 6.071 ± 0.935 | S1: 12.332 ± 1.526<br>S2: 10.763 ± 0.985<br>S3: 8.763 ± 0.880 | S1: 5.097 ± 0.764<br>S2: 5.561 ± 0.691<br>S3: 5.988 ± 0.605 |
| Inter-ch3ain<br>average COM<br>distance | S1: 1.947 ± 0.134 | S1: 1.781 ± 0.049 | S1: 1.698 ± 0.157 | S1: 1.726 ± 0.062 |
|  | S2: 1.847 ± 0.132 | S2: 1.854 ± 0.059 | S2: 1.599 ± 0.101 | S2: 1.676 ± 0.075 |
|  | S3: 1.725 ± 0.137 | S3: 1.818 ± 0.070 | S3: 1.889 ± 0.104 | S3: 1.700 ± 0.052 |

The C-terminal hook model^24^ suggests salt-bridges between Lys-24^a^- Asp75^b^ and Asp76^b^-Lys71^b^ in having a critical role in the dimerization of HGI-III. Asp75and Asp76 are part of the C-terminal tetra-peptide sequence which is missing in HGGI isoforms; its absence attributed to the existence of HGGI as a monomer. Contrary to the model, present simulations on HGI_1-76-FL consistently suggest that there is no stable salt-bridge (SB) interaction between Asp75^b^/Asp76^b^ and Lys24^a^/Lys71^b^. The starting protein structure for full length HGI simulations was modelled to emulate the C-terminal hook and the proposed salt-bridge interactions; a complete destabilization of the C-terminal hook is observed in the simulations (Figure 4a). Instead, the Lys24 residue forms a relatively stable salt bridge with Asp18 of the opposite subunit. The plots in Figure 4b comparing stability of these SB interactions, distinctly show the formation of Lys24-Asp18 SB at both ends of the dimer interface. Thus, it may be a significant factor in the dimerization of HGI-1_76. However, it fails to justify the experimentally observed monomeric existence of HGGI-I, II and III. Since neither Asp18 nor Lys24 is missing in these truncated isoforms, they should also exist as dimers. This inconsistency indicates that there may be additional factors that influence the homodimer formation.

**Figure 3.**
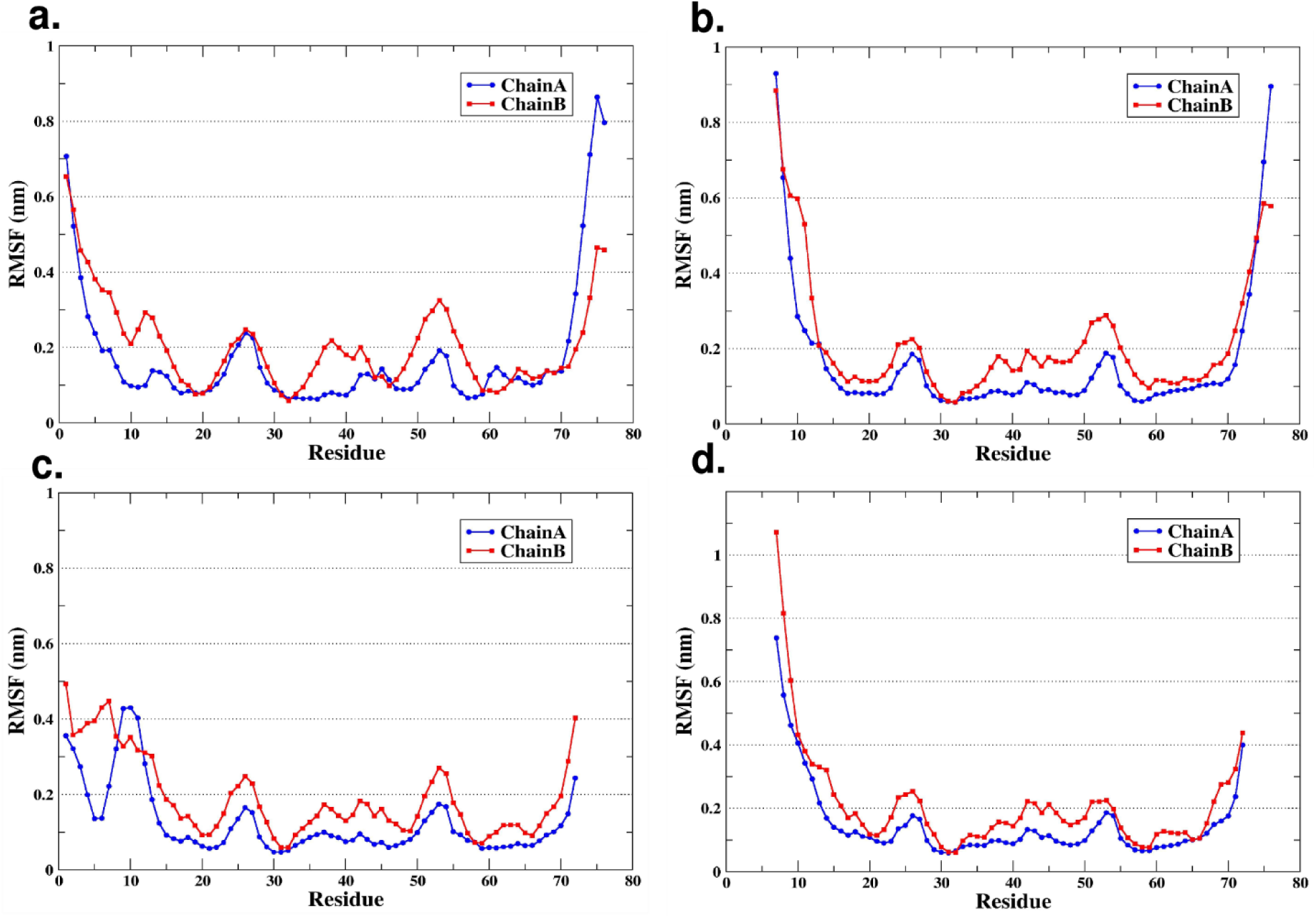
RMSF plots of residue Cα atoms in case of a) HGI_1-76-FL and its truncated forms b) HGI_1-72, c) HGI_7-76 and d) HGI_7-72. The blue and red plots depict RMSF for individual subunits.

**Fig 4:**
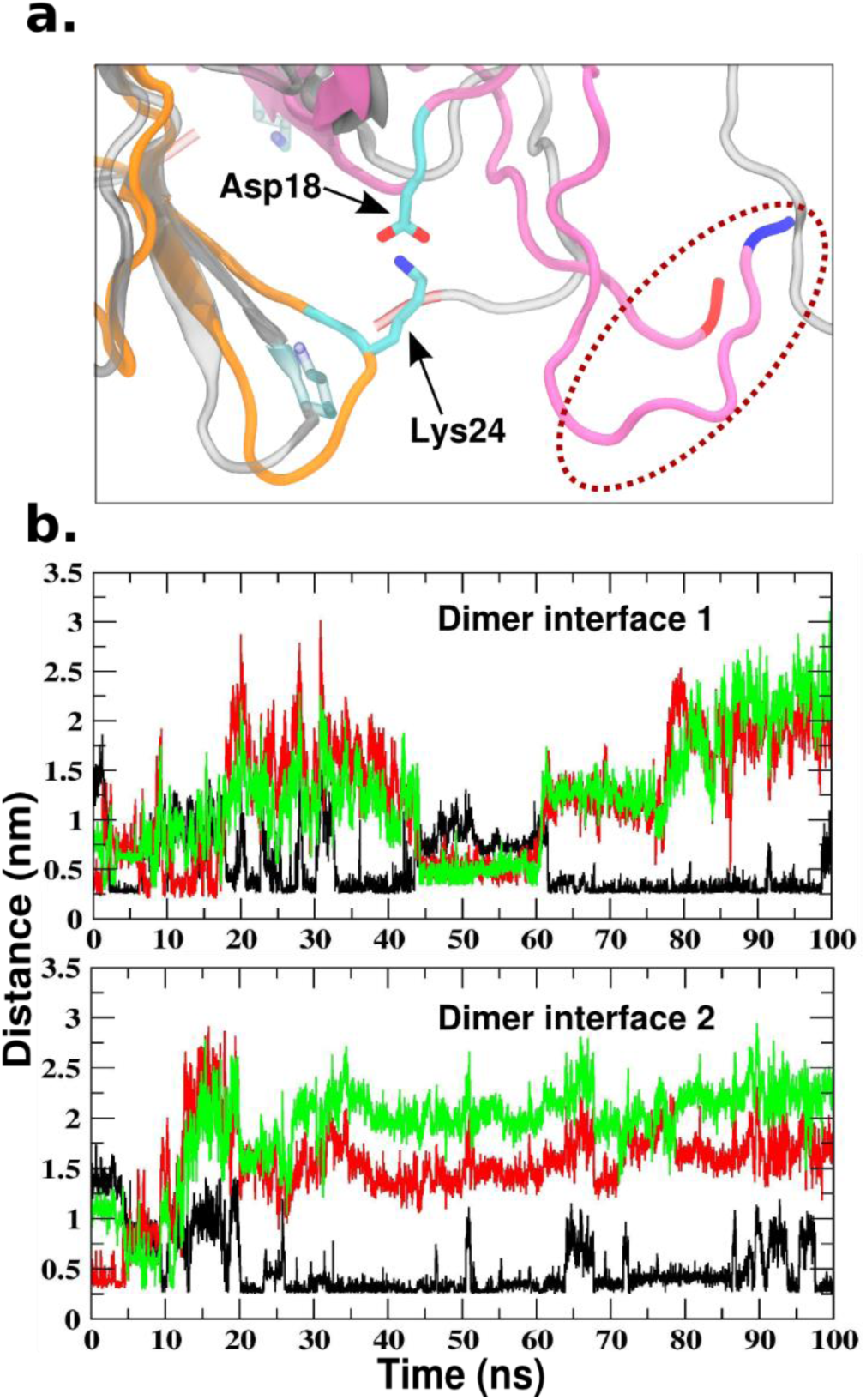
a) Superposition of initial and final coordinate of HGI_1-76-FL trajectory showing destabilization of the C-terminal hook and formation of salt-bridge between Lys24 and Asp18. Initial coordinates are depicted in translucent Gray showing the C-terminal hook conformation. Final structure shows salt-bridge between Lys24 and Asp18 (in Cyan) from opposite HGI monomers. C-terminal is destabilized and interacts with the N-terminal of the same chain. C and N terminals are marked as Red and Blue respectively. b) Plots comparing distance between Lys24 and Asp18 (black), Asp75 (green) and Asp76 (red) indicating greater stability of SB interactions along the full 100ns trajectory. The two plots represent the two ends of dimer interface with monomers in a two-fold symmetry. Lys24-Asp18 SB has a greater stability along the full trajectory.

Analysis of the simulations on HGI_1-72, HGI_7-76 and HGI_7-72 was similarly done to assess behavior of Asp18 and Lys24 in these systems. It should be noted that HGI_7-72 system is equivalent to the truncated form of HGI i.e. HGGI-I found in the germinated Horse Gram seeds. As expected, Lys24 forms a salt bridge interaction with Asp18 in all simulations, regardless of the presence/absence of the N or C terminal ends. One particularly interesting characteristic of this salt bridge interaction is the intermittent disruptions (for timespans as long as 4-5 ns) along the length of almost all the simulations (Figure 5). Although salt bridges generally undergo thermal fluctuations and continuously undergo breakage and reformation of bonds, the extent of the such disruptions (in other words the strength) depend on a number of factors such as charge on interacting ion pairs, presence of other ionic groups in vicinity, flexibility of the backbone and whether the salt bridge is buried/solvent exposed.^49,50^ Accordingly, Asp-Lys salt bridges would be considered the weakest of all salt bridges, more so, when they are exposed to aqueous solvent or in close vicinity of competing charged residues.

**Figure 5:**
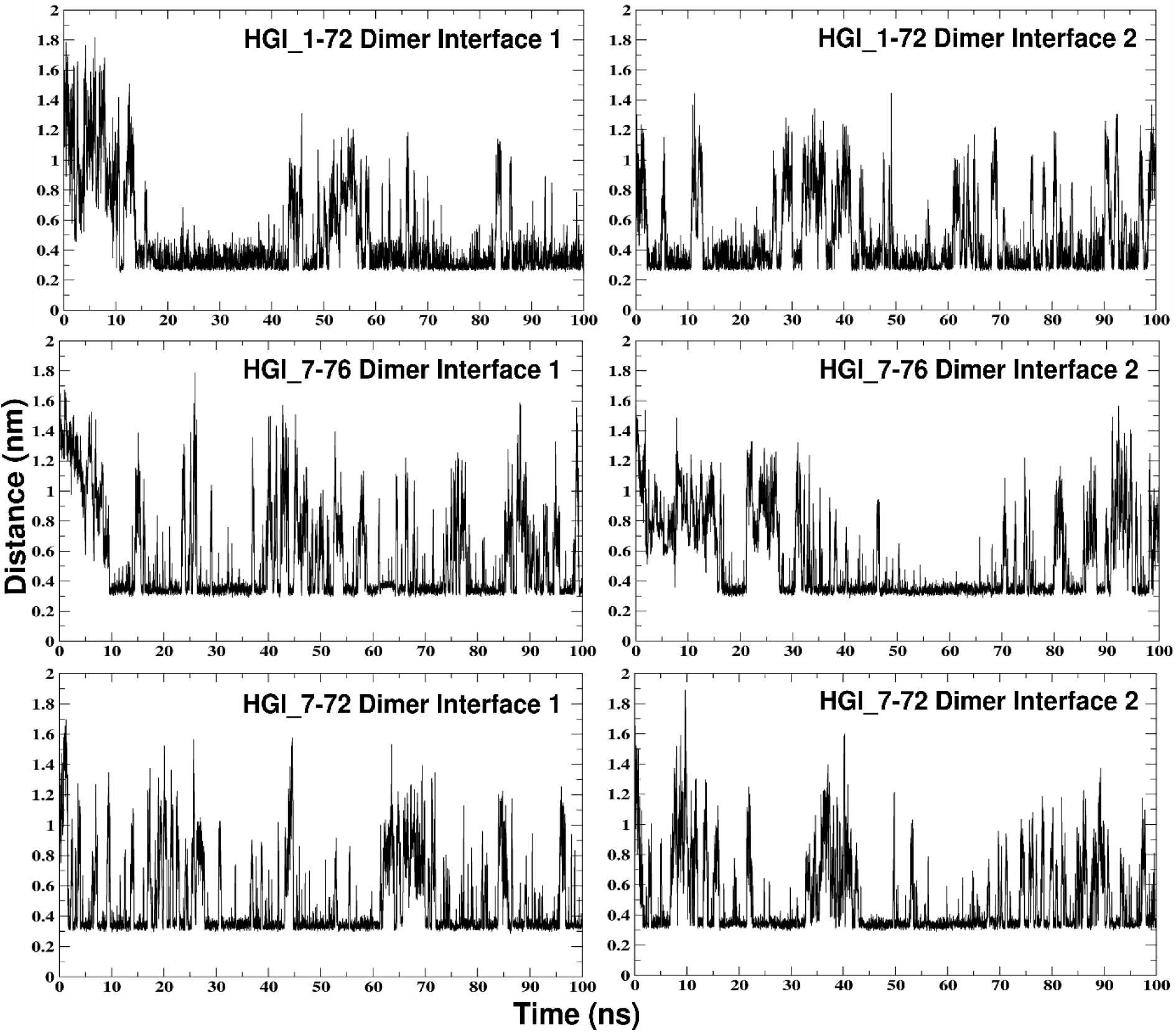
Stability of Lys24-Asp18 salt bridge interaction in simulation trajectories of HGI_1-72, HGI-7_76 and HGI-7_72.

Since the residues in question are surface residues, solvent exposure could be a possible factor influencing the interactions. Thus, the relative solvent accessibility of the amino acids Lys24^a^, Asp18^b^, Asp75^b^ and Asp76^b^ was analyzed. Figure 6 shows the relative solvent accessibilities of the four amino acids for coordinates extracted from a single simulation trajectory of HGI-1_76. The two graphs represent the two diametrically opposite ends of HGI-III. The general trend deduced from the plots is the relatively high solvent accessibility of Lys24 (>50%). The solvent accessibilities of both Asp75 and Asp 76 are much greater (∼75%) compared to Asp18 (∼30%). Moreover, Asp18 has a greater backbone stability compared to Asp75 and Asp76 (Figure 2a). These observations together provide a possible explanation as to why Lys24 fails to maintain a stable salt bridge interaction with Asp75/Asp76 and instead seeks to form interaction with Asp18. Figure 6 depicts a significant decrease in the solvent accessibilities of Asp75/Asp76 after 20 ns. Visual analysis of the trajectories revealed that this was due to the interaction of the C-terminal end with the N-terminal end. Similar trends were obtained from analysis of additional independent simulations of HGI-1_76.

**Figure 6:**
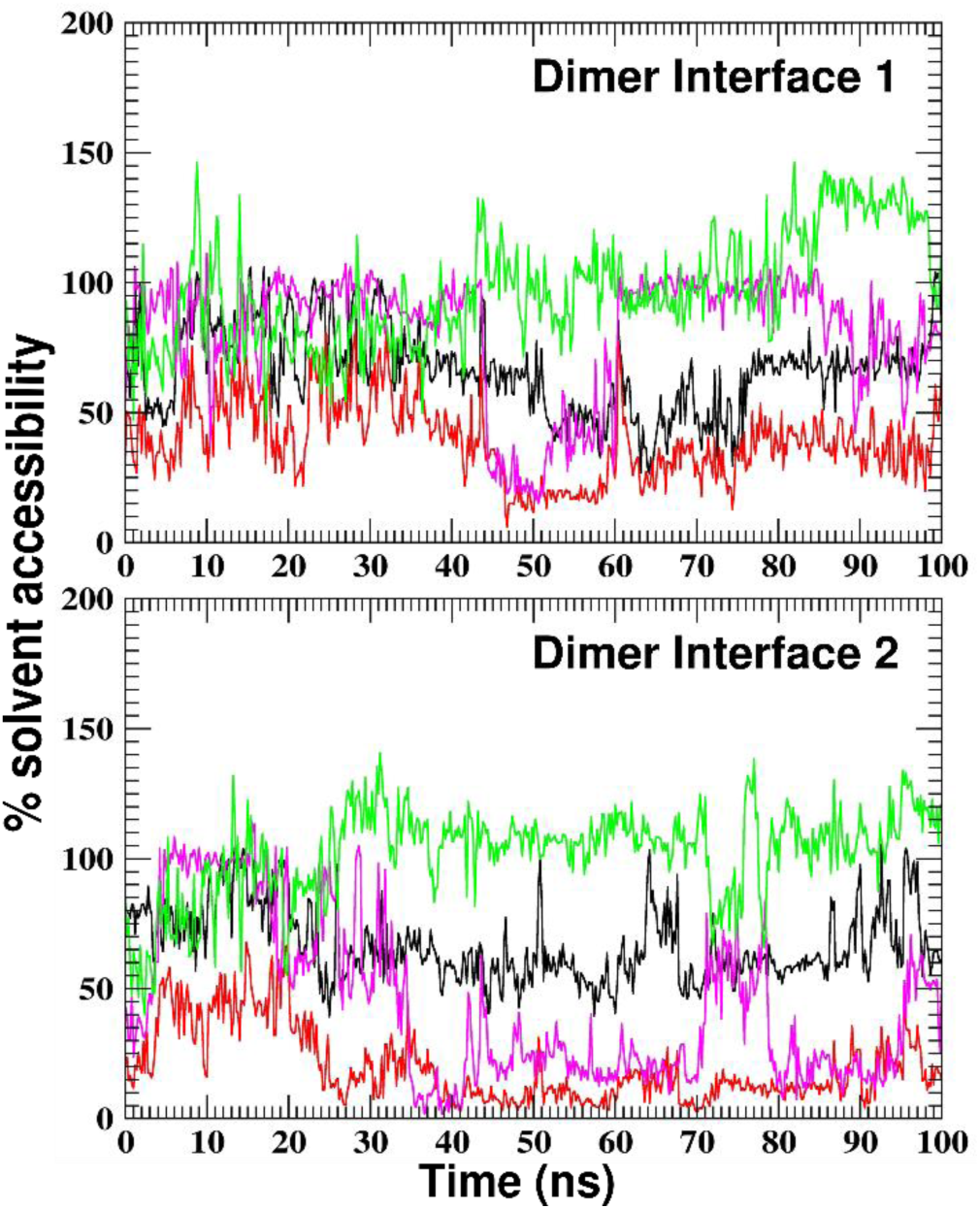
Solvent accessibility of residues Lys24 (Black), Asp18 (Red), Asp75 (Magenta) and Asp76 (Green) in HGI-1_76 simulation. Two plots represent the opposite ends of the dimer interface.

To summarize, these results allow us to speculate that although Lys24^a^ and Asp18^b^ form a SB interaction, it may not be sufficiently strong to hold the dimer complex together; the C-terminal and/or N-terminal region (alone or in conjunction) could be the key elements that provide additional electrostatic interactions at both ends to hold the dimer in case of the full-length HGI. Another possibility is that the terminal regions could be acting together to shield the interfacial Lys24-Asp18 SB at the two ends from aqueous solvent thereby significantly increasing the strength of SB interaction. It is important to take note that in all simulations, the protein converges to configurations where the N-terminal interacts with the C-terminal of the same subunit (Figure 4a). Hypothetically, there could also be a scenario where the N-terminal of one subunit interacts with the C-Terminal of the second subunit, thereby holding the dimer. Structural analysis indicates a disulfide linkage between Cys16^a^ and Cys70^a^ which puts physical constraints on N-terminal to remain at one end, disallowing movement to diametrically opposite end.

### REST2 simulations for Improved Sampling of Phase Space

In previous section, the analysis of N and C terminal regions of dimeric HGI_1-76-FL showed that free N-terminal region changes from an extended form to a folded form that interacts with the C-terminal region. However, among all simulations, there was clear lack of a characteristic pose in which the terminal regions interact. Moreover, considering their flexibility the terminals may exist in an ensemble of states, each state with its own relative stability and probability of occurrence. It is hence reasonable to assume that the lack of specific binding pose(s) could be due to inefficient sampling of the conformational space. One of the drawback of conventional simulation methods is the inability of proteins to access the minimum energy conformations that are separated by high energy barriers.^51^ This results in system that is trapped in one of the numerous local energy minima spread throughout the energy landscape, the corresponding state significantly different from the native state. In order to obtain proper canonical distribution for a protein structure, efficient sampling methods have been developed and successfully employed in studies of disordered regions and protein folding.^25,51–55^

#### Full length HGI-III

To obtain the most probable interaction pose(s) for the disordered N and C terminal of full length HGI dimer, REST2^25^ method was employed (See Methods for details). For the 32 replicas with λ value ranging from 1.0 to 0.5 (effective temperatures from 300 to 600 K), an acceptance rate of 33-42 % was obtained. The backbone RMSD plot of the dimer at the lowest λ replica indicates better sampling efficiency as compared to classical MD at the same temperature (Figure 7a, red plot compared with Figure 2a, blue plot). Also, the increasing RMSD value implies that the there is a gradual departure from the initial state while the protein samples the most visited conformations. A maximum RMSD value of about 1.3 nm was observed relative to the initial configuration. However, the backbone RMSD plot of the dimer discounting the flexible terminal segments is stable around 0.2 nm suggesting stability of the core structure (Figure 7a, black plots). Also, the secondary structural elements (chiefly Beta strands and turns) in the dimer are maintained throughout the simulation (Figure 7b). Hence, it is evident that at 300 K, the sampled states of the protein differ largely in the conformation of most disordered loops or random coils i.e. the free N and C terminal region. Although, the protein is essentially devoid of any alpha helix structure, there is transient formation of alpha helix (in blue) containing 3-4 residues as the protein samples different conformational states. These are formed in the terminal region in due course of sampling as confirmed by visual inspection of the trajectory. Since no sequence specific association or structural stabilization could be inferred based on the position of the helix, it was not considered as functionally relevant.

**Figure 7.**
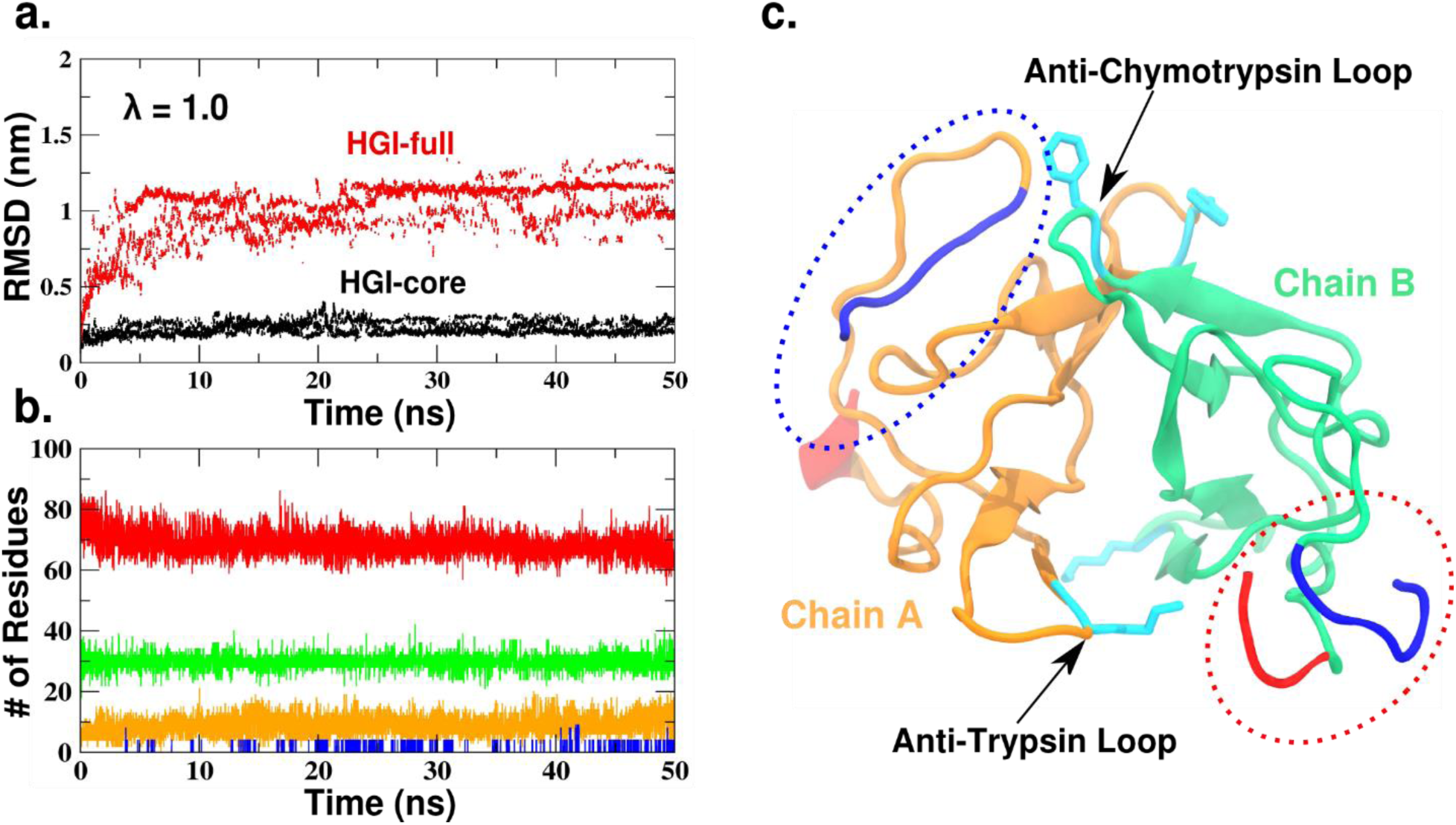
REST2 results for HGI-III. a) Graph depicting RMSD values of lowest replica frames (λ=1.0) obtained by REST2 simulations of HGI-III. The red plots denote the RMSD of the full protein (1-76) and black represents the RMSD of core residues (16-70). b) Secondary structure content of the sampled configurations. Red, Green, Yellow and Blue plots denote Coils, Beta-strand, Turns and Alpha-helix respectively. c) Representative coordinate for the topmost cluster obtained from cluster analysis of REST2 data. The terminal regions are colored in Red (C-terminal) and Blue (N-terminal). The dotted red ellipse marks the Anti-Trypsin Loop Proximal (ATLP) configuration formed by terminals of Chain B (in green), that can be seen close to the Anti-Trypsin loop of the opposite monomer, Chain A (in orange). The dotted blue ellipse marks the Anti-Chymotrypsin Loop Proximal (ACLP) state assumed by the terminal of Chain A. N-terminal is stabilized in a position close to the Anti-Chymotrypsin loop and C-terminal is in a freely suspended state.

Cluster analysis of the trajectory corresponding to 300 K (λ=1.0) was conducted to identify the most visited configurations at the temperature during the course of the simulation. Since the full length HGI has a significantly long, flexible terminal region, a relaxed RMSD cut-off of 0.25 nm was used for the analysis. A total of 265 clusters were identified in the analysis which is again indicative of the large conformational space sampled by the protein. A plot depicting cluster number of each time frame in the trajectory reveals that the cluster configurations are spread along the full trajectory (Figure 8). A relatively tight clustering is seen in case of the first 5 clusters compared to the rest. The top 20 cluster representatives were visually analyzed for identifying interesting configurations which may be biologically relevant. The first cluster which accounts for 27% of all the sampled configurations is shown in Figure 7c. The highlight of this configuration is the interaction between N and C terminal ends of one monomer (Chain B) forming a structural ‘appendage’ which lies in proximity to the anti-trypsin loop of other monomer (Chain A). Preliminary structural analysis showed that this structure is stabilized by 12 hydrogen bonds (Table 3) and a salt-bridge between Arg36 and Asp75. Out of these, 4 hydrogen bonds are exclusively formed between the N and C terminal segments that form the basic framework of this structure, whereas the remaining interactions seem to stabilize the structure relative to the rest of the chain. On the other hand, the N and C terminal regions of Chain A are seen to assume an open configuration, where the N-terminal region forms a wide hairpin-like loop which is close to the two anti-chymotrypsin loops. The N-terminal hairpin is stabilized in this position by as many as 7 hydrogen bonds (Table 4) and salt-bridge interaction between Glu11 and Arg36. No interactions with the Chymotrypsin loops were detected. For purpose of further discussion, hereafter the two forms of configurations will be termed as anti-trypsin loop proximal (ATLP) and anti-chymotrypsin loop proximal (ACLP) state. Interestingly, out of top 5 cluster (∼50% of all configurations), configurations in clusters 3 and 4 closely (each ∼5% of all configurations) resemble that of the top cluster (Figure 8b). Taken together, this configuration as described above accounts for about 37% of all the sampled states. This is a significant proportion considering the high flexibility of the terminal regions. The proximity of these prominent configurations to the two reactive loops in HGI suggests a functional association with trypsin and chymotrypsin binding.

**Table 3.** Hydrogen Bonds stabilizing the Anti-Trypsin Loop Proximal (ATLP) configuration. Bonds involved in N and C-terminal backbone stabilization marked by (*). # denotes bonds that remain stable in subsequent 50 ns simulation.

| Donor | Acceptor | Donor Atom | Acceptor Atom | H-bond Distance (Å) |
| --- | --- | --- | --- | --- |
| His2<sup>#</sup> | Gln4 | NE2 | OE1 | 2.910 |
| Ser12<sup>#</sup> | Pro9 | N | O | 3.396 |
| Lys14<sup>#</sup> | Asp76 | NZ | OC2 | 2.795 |
| Arg36<sup>#</sup> | Asp76 | NH1 | OC1 | 2.878 |
| Asn38<sup>#</sup> | Glu11 | ND2 | OE2 | 2.943 |
| Ser42 | Lys14 | N | O | 2.938 |
| Ser42<sup>#</sup> | Lys14 | OG | O | 2.797 |
| Lys71* | Asp1 | N | OD2 | 2.806 |
| Ser72* | Asp1 | N | OD1 | 3.106 |
| Ser72*<sup>#</sup> | Asp1 | OG | OD1 | 2.684 |
| Thr6*<sup>#</sup> | His74 | N | O | 3.282 |
| Lys14*<sup>#</sup> | Cys40 | N | O | 2.793 |

**Table 4.** Hydrogen Bonds stabilizing the Anti-Chymotrypsin Loop Proximal (ACLP) configuration.

| Donor | Acceptor | Donor Atom | Acceptor Atom | H-bond Distance (Å) |
| --- | --- | --- | --- | --- |
| His3 | Cys40 | NE2 | O | 2.918 |
| Ser13 | Glu11 | N | OE2 | 2.901 |
| Ser13 | Glu11 | OG | OE2 | 2.684 |
| Asn38 | Glu8 | ND2 | OE1 | 3.532 |
| Asn38 | Glu8 | ND2 | OE2 | 3.194 |
| Ser45 | Asp1 | OG | OD1 | 2.629 |

**Figure 8:**
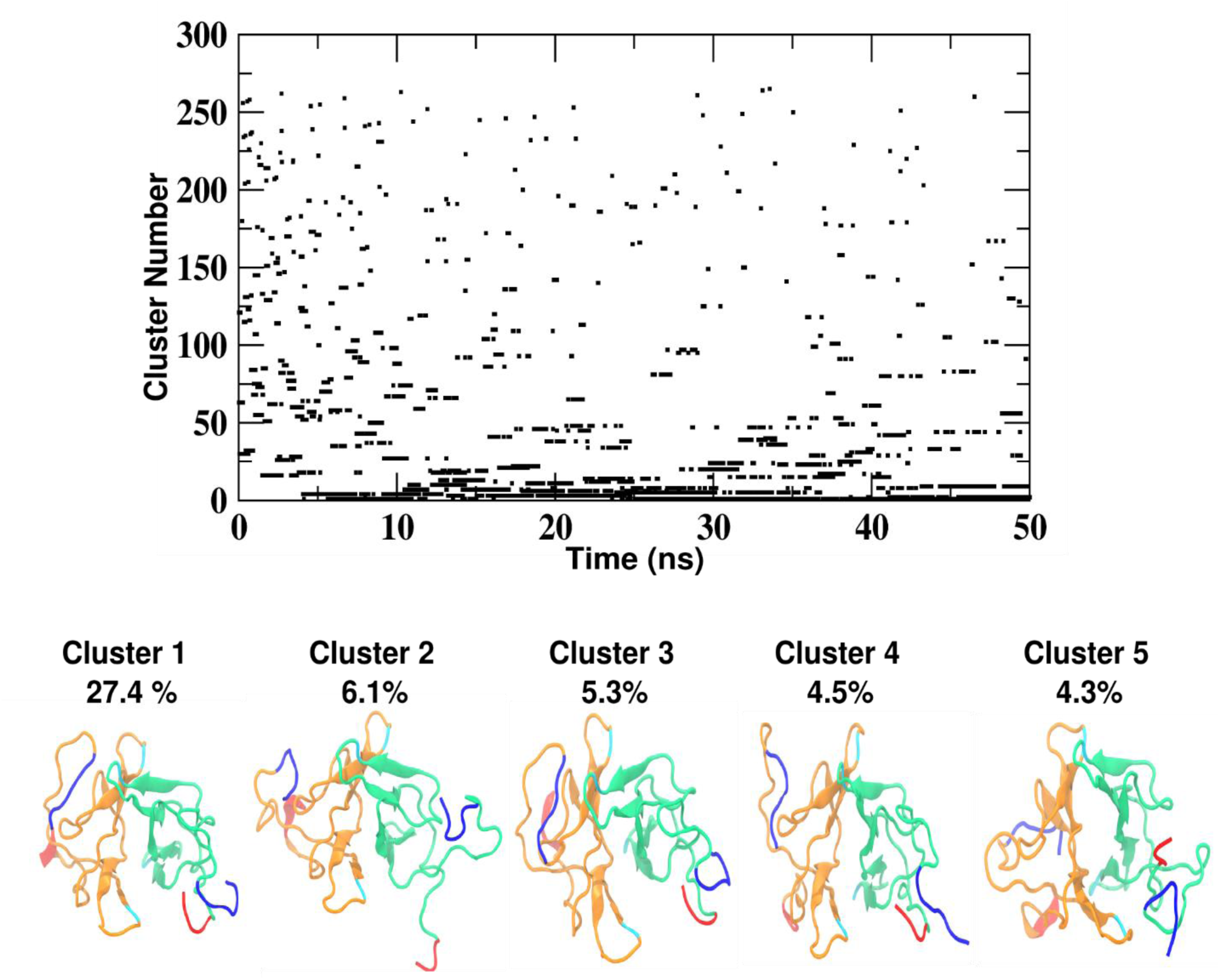
Cluster distribution of trajectory for the first replica (λ=1.0) from REST2 simulations of HGI-III. The top five cluster representatives are depicted along with their respective population percentages out of the total sampled configurations. The blue and red segments represent the N and C terminals respectively.

#### Truncated HGGI-I

The REST2 method was also employed to identify the significant configurations in case of HGI_7-72 (or HGGI-I). A comparison of the results with that of full length HGI could shed light on the differences in the conformational states available to the two proteins and how these differences lead to the observed behavior i.e. dimerization in case of full length whereas HGGI-I exist as monomers. An acceptance rate of 41-48 % was achieved for the 32 replicas. Backbone RMSD plots for first replica (λ=1.0) indicated better sampling efficiency compared to classical MD simulations of HGGI-I (Figure 9a, red plots compared with Figure 2d). Here again, it is evident that the sampled structures chiefly vary in the flexible terminal region; RMSD plot of the core protein is stable around 0.2 nm (Figure 9a black plots). Since, the flexible terminal region includes just the N-terminal 7-15 residues, we see a maximum divergence of only 0.6 nm RMSD from the initial structure. Cluster analysis resulted in 286 clusters using a RMSD cut-off of 0.15 nm that are distributed along the full trajectory (Figure 10). Unlike HGI, the top clusters in this case were not seen to be highly populated with an relatively even distribution of the possible states (Figure 9c). The top cluster representative corresponding to 9% of sampled configurations is depicted in Figure 9d. Visual analysis of representative structures of the top 20 clusters showed that the truncated terminal regions of HGGI-I do not have any particular interaction with each other. The top configurations obtained have essentially open dimer interface with the Aps18-Lys24 SB exposed. But complete dimer disruption were not observed at 300 K in the course of present simulations. Interestingly, the N terminal region is seen to form helical turns in all of the top 5 cluster representatives and in 14 out of the top 20. The helix is formed specifically by the residues ^(10)^SESSK^(14)^ flanked by Proline residues at both ends. This is also evident from the secondary structure plot which indicates the formation of helix throughout the simulation (Figure 9b). In a previous study^23^ on HGGI forms, it was suggested that the N-terminal segment in HGGI isoforms (7-13 residues) has a negative effect on trypsin binding and inhibition (compare N-terminal sequences and K_i_s values of HGGI forms marked as * in Table 1). Whereas in HGI forms, the extra N-terminal segment and its length (1-7 residues) seems to positively correlate with trypsin inhibitory constant (compare N-terminal sequences and K_i_s values of HGI forms marked as # in Table 1). This suggests that after the proteolysis of the first six N-terminal residues in HGI forms, the remaining residues assume somewhat altered function in HGGI’s. It has been speculated that since residues that make up the N-terminal are predominantly negatively charged, electrostatic effects may be responsible for the observed variation in Trypsin inhibitory constants of HGGI’s.^23^ However, trypsin surface around the active site pocket has predominantly hydrophobic and basic patches.^56^ Hence, it is likely that the hydrophobic amino acids on trypsin surface are responsible for the decreased trypsin affinity in case of HGGI-I isoform.

**Figure 9:**
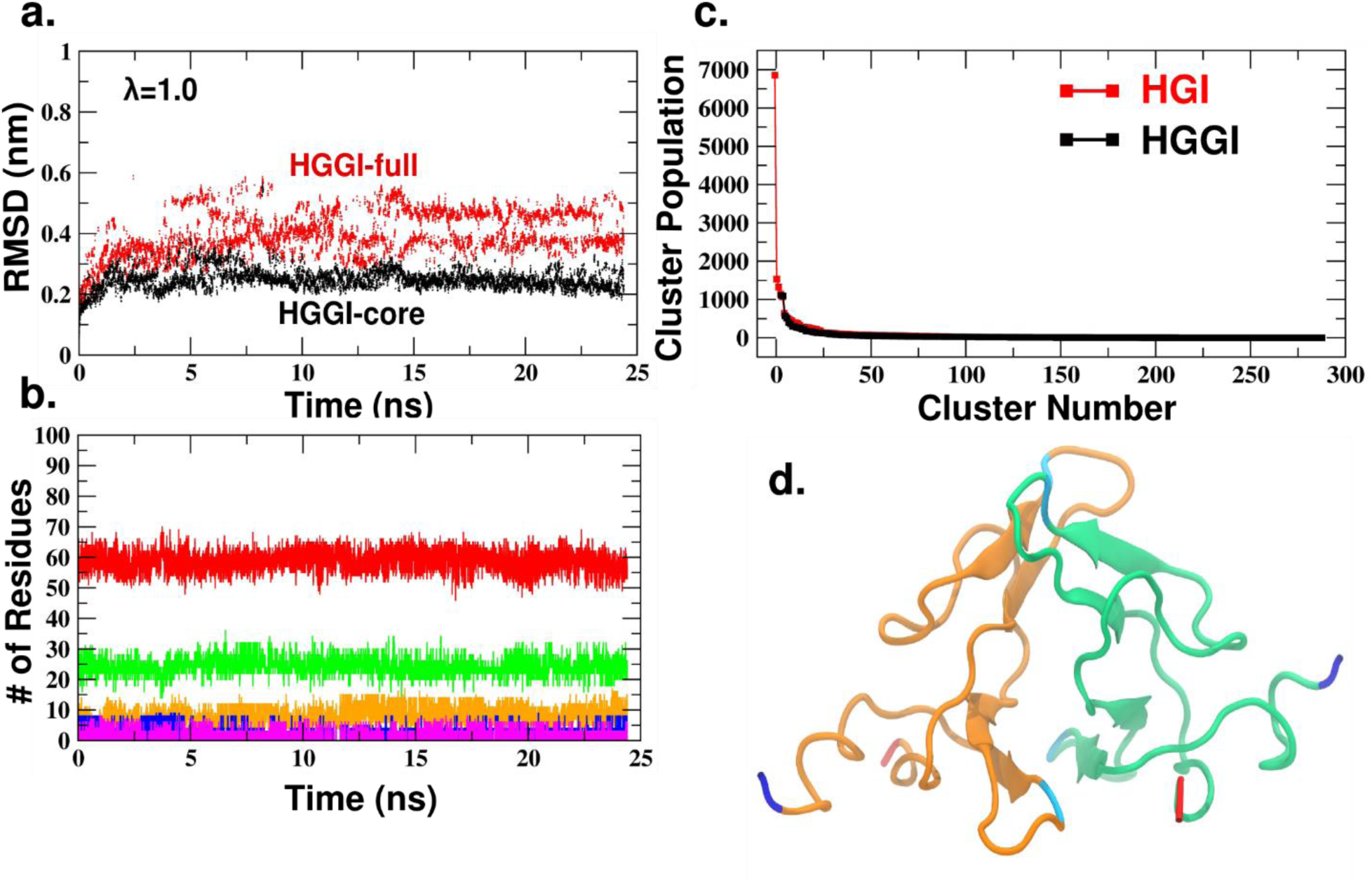
REST2 results for HGGI-I. a) Graph depicting RMSD values of lowest replica frames (λ=1.0) obtained by REST2 simulations of HGGI-I. The red plots denote the RMSD of the full protein (7_72) and black represents the RMSD of core residues (16-70). b) Secondary structure content of the sampled configurations. Red, Green, Yellow, Blue and Magenta plots denote Coils, Beta-strand, Turns, Alpha-helix and 310 Helix respectively. c) Comparison of plots representing cluster populations of HGI-III and HGGI-I. d) Representative coordinate for the topmost cluster obtained from cluster analysis of REST2 data. The terminal regions are colored in Red (C-terminal) and Blue (N-terminal). Helical conformation is seen in both N-terminals.

REST2 method employed here allowed enhanced sampling of the phase space and provided prominent structures that are visited more often by the full length and truncated form of the HGI. The top configurations obtained in each case can now be studied further to ascertain their stability. It is expected that configurations of the terminal region, as observed in case of full length HGI, would be relatively stable considering the presence of a number of stabilizing interactions. Notably, ATLP state wherein the N and C terminals interact in a specific manner to form extra folds concurs with the previously expressed possibility where such a structure may act to stabilize the dimer by supplying additional interactions and shielding of the Asp18-Lys24 SB from aqueous solvent molecules. This configuration utilizing a salt-bridge (Arg36-Asp75) and hydrogen bonds seems to provide the desired steady interface for binding the second subunit. The lack of the terminal segments would result in the absence of this stabilizing interface and a concomitant monomeric existence. This hypothetical model of dimer stabilization may explain the existence of HGI and HGGI forms as dimer and monomer respectively. The ACLP state was similarly not anticipated from the classical MD simulations. It is quite possible that such a configuration of N-terminal could have a functional role in Chymotrypsin binding considering its proximity to the Anti-Chymotrypsin Loop.

### Molecular Dynamics of HGI configuration

A somewhat static picture of the configurations obtained from the REST2 simulations allowed limited analysis and understanding of the prominent protein models. In order to obtain better insight into the stability of the prominent configurations of N and C terminals, classical MD simulation was performed on the top cluster representative (as depicted in Figure 7c). The RMSD calculated over the 50 ns trajectory indicates a greater stability as there is little divergence from the input state (Figure 11a red plot). Slight deviation observed is attributed to the terminal regions as the RMSD of core domains of HGI remain stable throughout the simulation (Figure 11a black plot). The backbone flexibility of individual residues of each monomer chains is represented as RMSF plots in Figure 11b. Backbone of terminal regions of Chain A which are in the ACLP state show high mean fluctuations (black plots). The N terminal of Chain A is stable to an extent but has greater flexibility around the residues 6-8, a tripeptide segment Thr-Asp-Glu. This region is part of N-terminal which is nearest to the anti-chymotrypsin loop. On the other hand, the red plot shows that both terminal region of Chain B are quite stable. This is because the Chain B terminal regions exist in the ATLP state wherein they interact with each other via multiple hydrogen bond interactions. The only significant backbone fluctuation in Chain B is at the N terminal Asp1 residue.

**Figure 10:**
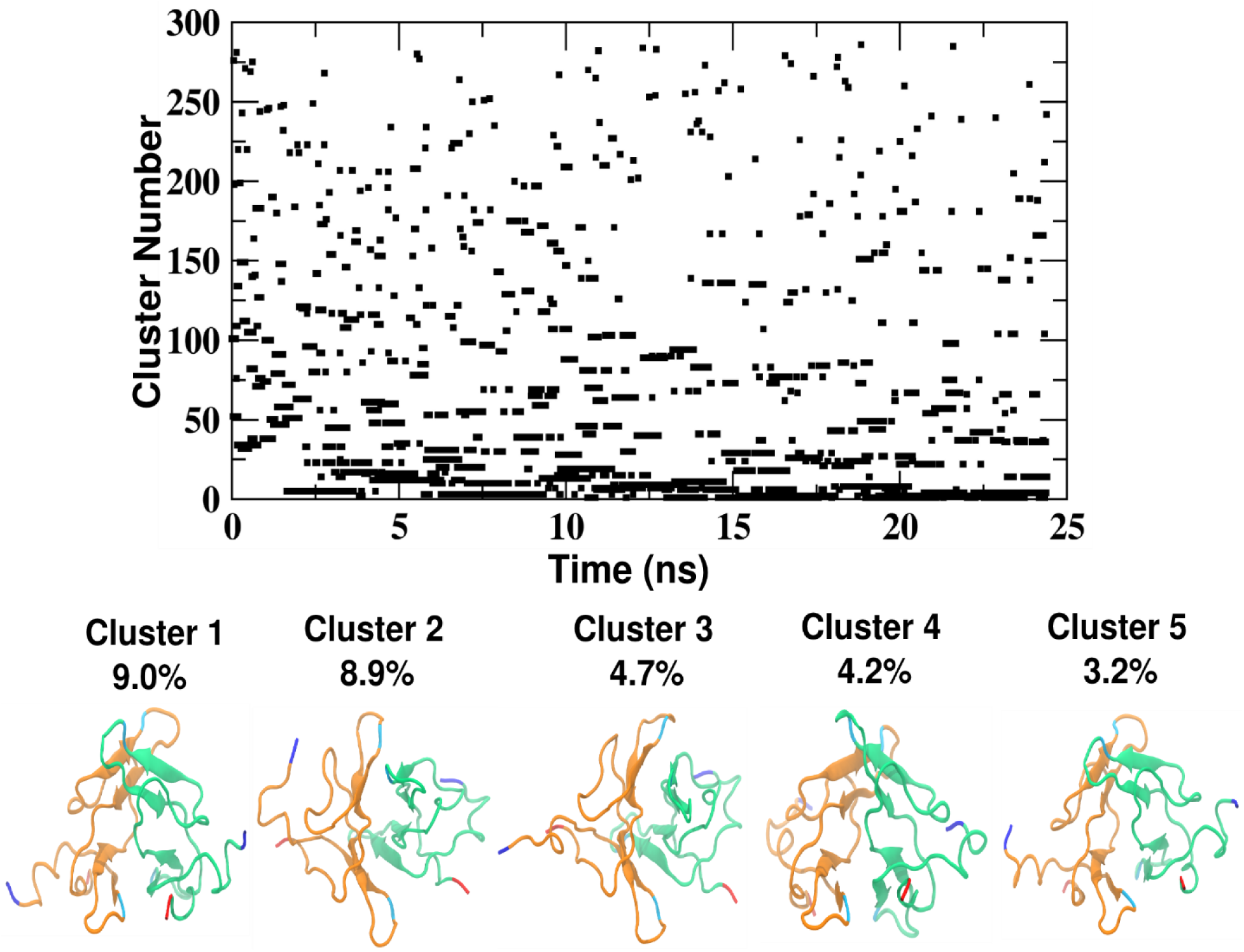
Cluster distribution for trajectory corresponding to first replica (λ=1.0) from REST2 simulations of HGGI-I. The top five cluster representatives are depicted along with their respective population percentages out of the total sampled configurations. The blue and red segments represent the N and C terminals respectively.

**Figure 11:**
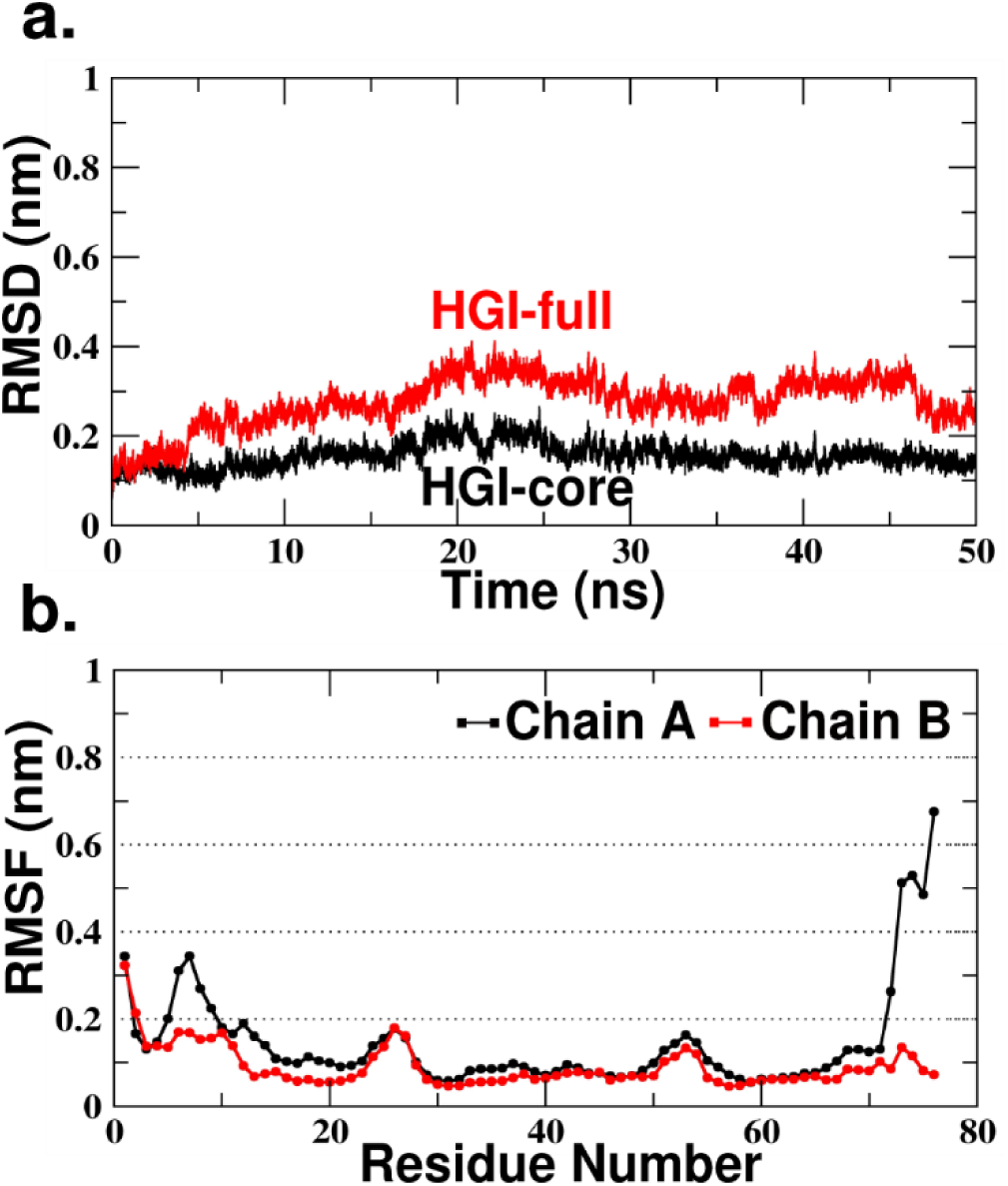
a) RMSD plot for 50 ns conventional MD simulation of the topmost cluster configuration suggesting a more stable configuration. Red and Black plots represent backbone RMSD for full-sequence (1-76) and core residues (16-70) of HGI-III respectively. b) RMSF plots for Cα atoms of each monomer chain.

As stated previously, the role of this ATLP structure in dimerization is of interest. Analysis of the simulation trajectory revealed that most of the hydrogen bonds holding ATLP configuration are stable throughout simulations (marked with *#* in Table 3). The salt-bridge between Arg36 and Asp75 is seen to be stable and hence critical to the stabilization of the C-terminal end in vicinity of the dimer interface (Figure 12a and 12b). Arg36 also forms hydrogen bond with the C-terminal carboxyl oxygen promoting greater stability. The Arginine side-chain is itself stabilized by interactions with neighboring Asp34 and backbone interactions with Cys57. Lys14 also forms a hydrogen bond with one of the C-terminal oxygen. The N and C terminal segments interact with each other in an anti-parallel arrangement involving 4 hydrogen bonds (Figure 12c). The N-terminal Asp1 which was stabilized by two hydrogen bonds in the initial frames is no longer involved in any interaction and suspends freely in the surrounding solvent. The surface electrostatics of ATLP region, illustrated in Figure 12d, shows the presence of a large negatively charged patch within the dimer interface portion formed by the ATLP structure. The Lys24 residue of the opposite monomer lies in close contact with this region. The inset in Figure 12d shows 4 Asp residues contributing to this patch. Lys24 which interacts chiefly with Asp18 also forms transient interactions with Asp75 side chain and backbone. Result from previous experimental studies^21,24^ showing monomeric existence of Asp75Ala, Asp76Ala and Lys71Ala mutants of HGI also corroborate with the present model. These residues form a crucial part of the assembly of terminal regions forming the ATLP structure, mostly stabilized by electrostatic interactions. It can be reasoned that any mutation to non-polar residues could significantly affect the interaction energy of the structure and lead to destabilization of the ATLP state. Together, the above observations and arguments lend support to the notion that both N and C-terminal ends play an intimate role in dimer stabilization by forming an additional interface region, that in addition to steric exclusion of solvent, also presents an interaction “hot-spot” on its surface for enhanced dimer stabilization.

**Figure 12:**
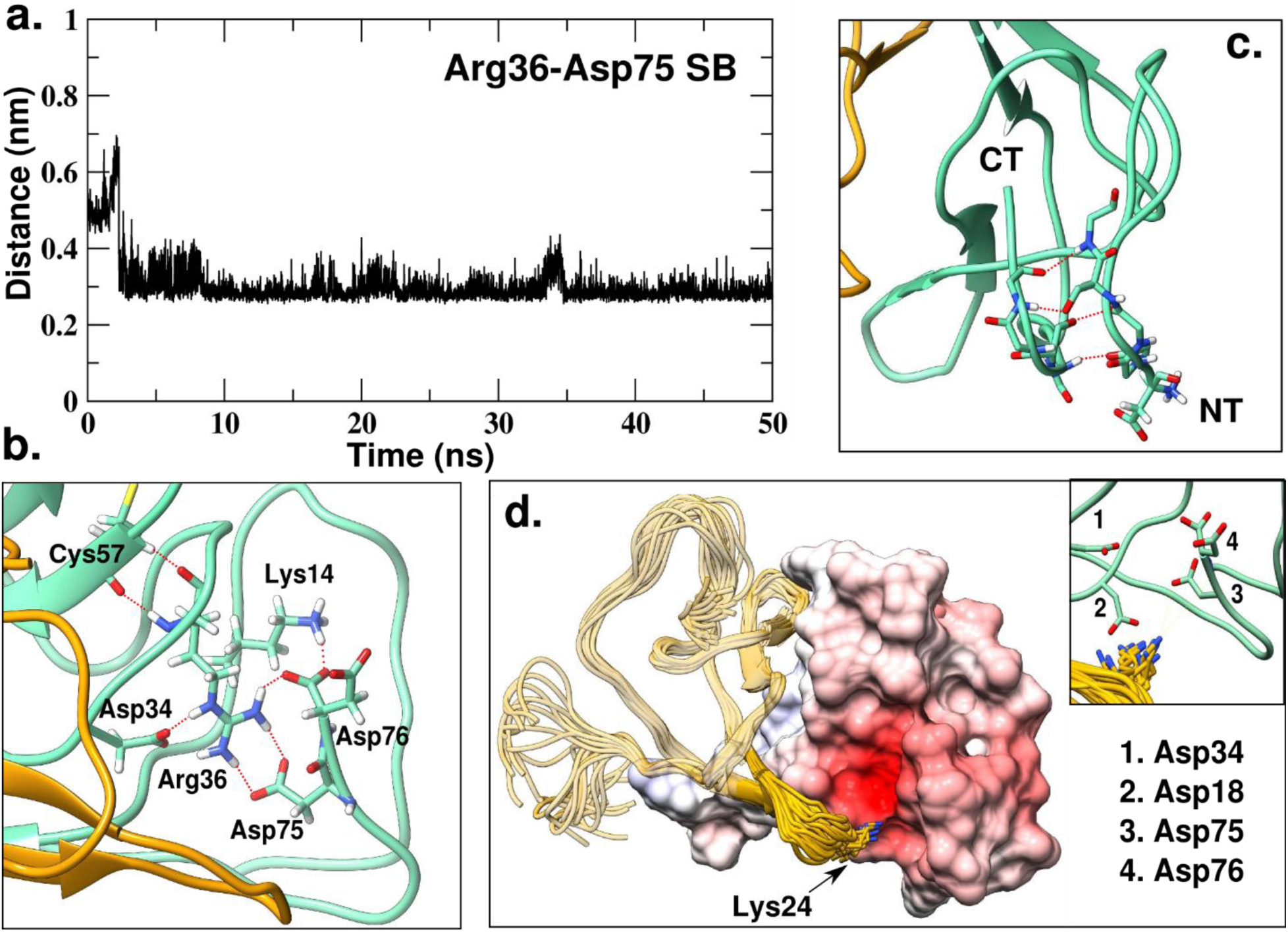
a) Plot depicting stability of Arg36-Asp75 salt-bridge belonging to the ATLP state. X-axis denotes distance between NH1 and OD1 of Arg36 and Asp75 respectively. b) Interactions involved in stabilization of C-terminal residues in the ATLP configuration. c) Interactions between N and C-terminal backbone in ATLP configuration. d) Surface Electrostatics representation of the HGI chain with terminals in ATLP state showing a negatively charged patch formed by the ATLP state. Electrostatics calculations were performed using APBS.^29^ Inset: Residues contributing to the negatively charged patch.

The protein system used for the current simulation affords the advantage of observing the effect of both presence and absence of the ATLP structure. One end of the dimer interface has ATLP configuration which can be assumed as the closed state, whereas in the opposite end the terminals exist in the ACLP state leaving the interface exposed. A comparison of the distance between the two subunits at both ends of the dimer interface showed, the open-end has a larger mean distance of 11.36 ± 1.2 Å between the monomers and experiences slightly greater destabilization compared to closed-end with mean distance of 10.91 ± 0.9 (Figure 13). Even though the difference seems insignificant, it can be argued that at a molecular level the open, exposed end may simply be more accessible to an external protein (such as trypsin) and slight destabilization of the interface and salt-bridge (Lys24-Asp18) disruption could significantly enhance accessibility to the Anti-Trypsin Loop. In comparison, the ATLP structural motif supplants a protective and stable cap which restricts access via steric effects, in addition to dimer stabilization via electrostatic interactions.

**Figure 13:**
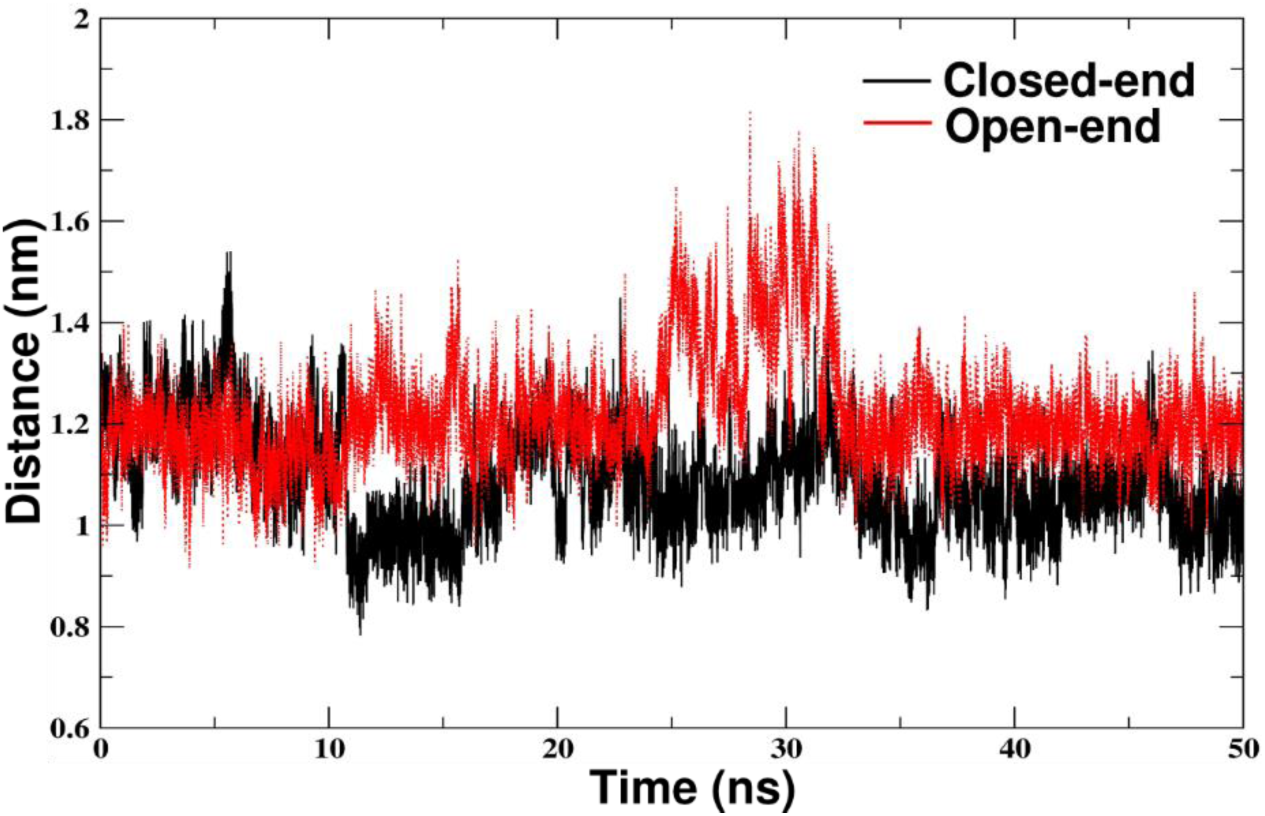
Comparison of the distance between the two subunits at the closed (ATLP) and open (ACLP) end of the dimer interface. Distance is calculated at both ends between backbone atoms in the B1 strand of each monomer.

Though, the above model explains the dimerization of HGI and its subsequent dissociation to monomers on proteolysis of the terminal segments, the mechanism of dimer to monomer transition still remains unexplained. The requirement of dimer disruption in HGI for trypsin/chymotrypsin binding has been shown by Kumar et. al.^24^ Based on intrinsic fluorescence quenching, they estimated the dissociation rate constants of dimer (HGI-III) and monomer (HGGI-I) for trypsin binding to be 5.92 μM and 5.45 μM respectively and for chymotrypsin to be 13.48 μM and 12.37 μM respectively. The stoichiometry was calculated to be 1:1 for both trypsin and chymotrypsin binding to the HGI-III dimer. In the present simulations, we have observed that 1) most prominent configurations of the terminal region i.e ATLP and ACLP are both are significantly stable and 2) the terminals of HGI exist in an ensemble of states at 300 K as evidenced by the large number of clusters obtained from sampling. Moreover, in subsequent simulation of the top configuration of HGI, we observed that the ATLP configuration is quite stable except for the N-terminal Asp residue. High backbone fluctuation of Asp1 together with its accessibility to external interactions makes it a likely point for initiation of a destabilization event. The trypsin inhibitory constants of HGI forms obtained from previous studies indicate that the binding to trypsin is linked to the N-terminal sequence (compare trypsin K_i_s values and sequences of HGIs in Table 1). Specifically, HGI-III which possesses the full amino-acid sequence has the highest binding affinity to trypsin (as indicated by the lowest K_i_s value). But, as the terminal residues are removed (in HGI-I and II forms) the affinity to trypsin decreases. Incidentally, inspection of the crystal structure of Cowpea BBI bound to trypsin showed the presence of a positively charged pocked in vicinity of the trypsin active site which might bind the N-terminal Asp. Taking these observations into perspective, we propose a model of dimer to monomer transition that is consistent with experimental observations. The dimer interface stabilized by the ATLP loop remains stable until the inhibitor interface comes in vicinity of trypsin. ATLP destabilization is initiated when the trypsin interacts with the exposed N-terminal Aspartate and changes the micro-environment around ATLP, sufficient to break interactions between N and C terminal. Once the N-terminal is destabilized, the C terminal assumes a disordered state and other ensemble states become more stable. The terminals gradually assume the ACLP configuration which simultaneously allows for dimer disruption and trypsin binding. The difference in the dissociation constant of dimer (5.92 μM) and monomer (5.45 μM)^24^ for trypsin implies a lag in trypsin binding to the dimer. This lag might be accounted by the time taken for destabilization of ATLP and transition to an alternate configuration. This mechanism also explains the decrease in trypsin binding affinity in HGI-I and II; first few N-terminal residues being absent does not allow trypsin to interact efficiently with the N-terminal end, thereby reducing the efficiency of dimer to monomer transition and affecting subsequent binding of trypsin.

## Conclusion

In summary, based on the present computational analysis, we have attempted to unify and explain the experimental results obtained regarding Horse Gram BBI forms. The biological requirement of different isoforms of HGI in dry and germinated seeds is not fully understood. The various forms have been known to have different characteristics such as inhibitory constants and dimerization preference, all seemingly dependent on the terminal sequences. Prior to our work, proposed models of dimerization have provided limited understanding of the dimerization process and factors involved in dimer to monomer transition. The involvement of N-terminal in dimerization and trypsin binding have not been factored previously. We have proposed a new model of HGI that agrees with the experimental observation more closely and describes the mechanism and factors that are involved in dimer/monomer formation and trypsin inhibition. The work is of immediate relevance to experimentalists and can aid in future studies on HGI to experimentally validate the model. Armed with understanding of such mechanisms, it is possible to implement the knowledge to the design of novel protein inhibitors.

## Acknowledgement

AA acknowledges Dr. Saravanan Murugeson and Harmanjit Singh for their gracious help with the computational work and proofing the initial drafts. Interdisciplinary Centre for Mathematical and Computational Modelling (ICM), Warsaw University, Poland is acknowledged for providing the computer facilities under Grant No. G55-11. AA thanks Prof. Balaji Prakash, CSIR-CFTRI, Mysore for his support and encouragement. Shasara Research Foundation is acknowledged for its financial support.

